# Effects of leg immobilization and recovery resistance training on skeletal muscle-molecular markers in previously resistance trained versus untrained adults

**DOI:** 10.1101/2024.07.12.603321

**Authors:** J. Max Michel, Joshua S. Godwin, Daniel L. Plotkin, Mason C. McIntosh, Madison L. Mattingly, Philip J. Agostinelli, Breanna J. Mueller, Derick A. Anglin, Alexander C. Berry, Marina Meyer Vega, Autumn A. Pipkin, Matt S. Stock, Zachary A. Graham, Harsimran S. Baweja, C. Brooks Mobley, Marcas M. Bamman, Michael D. Roberts

**Affiliations:** School of Kinesiology, Auburn University, Auburn, Alabama; Institute of Exercise Physiology and Rehabilitation Science, School of Kinesiology and Rehabilitation Sciences, University of Central Florida, Orlando, FL, USA; Healthspan, Resilience and Performance Research, Florida Institute for Human and Machine Cognition, Pensacola, FL, USA; Edward Via College of Osteopathic Medicine, Auburn, Alabama

**Keywords:** atrophy, hypertrophy, immobilization, bracing

## Abstract

We sought to examine how resistance training (RT) status in young healthy individuals, either well-trained (T, n=10 (8 males)) or untrained (UT, n=11 (8 males)), affected muscle size and molecular markers with leg immobilization followed by recovery RT. All participants underwent two weeks of left leg immobilization via the use of crutches and a locking leg brace. After this two-week period, all participants underwent eight weeks (3 d/week) of knee extensor focused progressive RT. Vastus lateralis (VL) ultrasound-derived thickness and muscle cross-sectional area were measured at baseline (PRE), immediately after disuse (MID), and after RT (POST) with VL muscle biopsies collected at these time points. T and UT presented lower ultrasound derived VL size (cross-sectional area and thickness) values at MID versus PRE (p≤0.001), and values increased in both groups from MID to POST (p<0.05); however, VL size increased from PRE to POST in UT only (p<0.001). Mean and type II myofiber cross-sectional area (fCSA) values demonstrated a main effect of time where PRE and POST were greater than MID (p<0.05) and main effect of training status where T was greater than UT (P≤0.012). In both groups, satellite cell number was not affected by leg immobilization but increased in response to RT (p≤0.014), with T being greater than UT across all time points (p=0.004). Additionally, ribosome content (total RNA) decreased (p=0.010) from PRE to MID while the endoplasmic reticulum stress proteins (BiP, Xbp1s, and CHOP) increased from MID to POST regardless of training status. Finally, the phosphorylation states of mechanistic target of rapamycin complex-1 signaling proteins were not significantly altered for either group throughout the intervention. In conclusion, immobilization-induced muscle atrophy and recovery RT hypertrophy outcomes are similar between UT and T participants, and the lack of molecular signature differences between groups supports these findings. However, these data are limited to younger adults undergoing non-complicated disuse. Thus, further investigation to determine the impact of training status on prolonged leg immobilization models mirroring current medical protocols (e.g., following orthopedic injury and surgery) is warranted.

## INTRODUCTION

Skeletal muscle is a highly plastic tissue that responds to various stimuli (1, 2, 3). Progressive resistance training (RT) is one of the most potent of these stimuli, with the hallmark adaptation to mechanical overload via RT including tissue and myofiber hypertrophy (4) whereas the disuse/unloading of the same muscle leads to a decrease in muscle size, or skeletal muscle atrophy (5, 6). There are many methods by which disuse-induced atrophy has been studied using human limb immobilization models such bracing or casting (7, 8, 9, 10, 11), spaceflight (12, 13, 14, 15), bedrest (16, 17, 18), and dry immersion (19, 20); all have proven to produce atrophy. While this phenomenon is somewhat well-studied, strategies proven to attenuate disuse-atrophy are less clear. One strategy to mitigate disuse-atrophy is the addition of intermittent high intensity muscle loading during a period of bedrest (17, 21). However, an injury or other event may be severe enough that physical conditioning during this time might become practically unfeasible. Therefore, attention has turned toward alternative parallel interventions that alleviate the disuse atrophy and/or treatments in series that expedite recovery upon reloading. The reader is referred to (5) and (6) for a more comprehensive review of mechanisms and prevention strategies of disuse atrophy, as well as recovery approaches. Briefly, while it is commonly thought that muscle protein synthesis (MPS) is the largest governing factor of disuse-atrophy, several molecular markers such as proteolytic proteins, ribosome content, and endoplasmic reticulum stress related proteins have been shown to be differentially regulated in response to disuse. While data indicating MPS declines with disuse are consistent, potential (dys)regulation of the translational machinery (i.e. ribosomal RNA, mTOR signaling) and proteolytic signaling pathways are not as well characterized in response to disuse-atrophy or subsequent RT-mediated regrowth. These markers were therefore selected for measurement as a part of this study.

It has been suggested that RT preconditioning could enhance RT adaptation after a period of training cessation, potentially by a “muscle memory” effect of muscle to RT adaptation (22, 23). This is not, however, the most applicable method for disuse-atrophy mitigation given that it would require the foresight of a disuse event and/or the ability or willingness to perform RT weeks to months prior to disuse. Whether habitual RT plays a significant role in the adaptation to and recovery from a disuse period has scarcely been investigated (24). Notably, as a lifestyle change, this is a modifiable factor without the foresight of an impending disuse event. Therefore, the purpose of this study was to examine if RT status affects neuromuscular function, skeletal muscle tissue, and cellular-level adaptations to 14 days of disuse and subsequent recovery via eight weeks of prescribed/supervised RT, were differentially affected by training status. We recruited 10 college-aged participants with 6±4 years of self-reported resistance training experience (T) and 11 untrained counterparts (UT) of similar age to engage in two weeks of unilateral leg immobilization immediately followed by eight weeks of bilateral knee extensor focused RT. Vastus lateralis (VL) ultrasound-derived thickness and muscle cross-sectional area were measured at baseline (PRE), after disuse (MID) and after eight weeks of RT (POST). VL muscle biopsies were also collected at these time points to examine the expression of atrophy and hypertrophy related proteins, total and ribosomal RNA content, satellite cell abundance, and fiber type specific myofiber cross-sectional area (fCSA). We hypothesized that T would experience more disuse atrophy than UT, but would present a more robust recovery from disuse (i.e., more hypertrophy) following eight weeks of RT. We additionally hypothesized that skeletal muscle markers of proteolysis and endoplasmic reticulum stress would present a signature aligned with these phenotypic responses.

## MATERIALS AND METHODS

### Participants and Ethical Approval

This study was conducted with prior review and approval from the Auburn University Institutional Review Board and in accordance with the most recent revisions of the *Declaration of Helsinki* (IRB approval no: 23-220 MR2305, clinical trial registration no: NCT05760066). The participants were young males and females from the local area that met the following criteria: (i) 18-35 years old; (ii) no known cardiometabolic disease (e.g. diabetes, hypertension, heart disease) or any musculoskeletal condition contraindicating participation in exercise training or donating muscle biopsies; (iii) free from metal implants that would interfere with x-ray based data collection; (iv) had not consumed known anabolic agents or agents that affect hormone status within the past two months (e.g. exogenous testosterone, growth hormone, etc.); (v) free from blood clotting disorders that would contraindicate donating a muscle biopsy; (vi) either no participation in a RT program more than 1 day/week for the previous year (UT group) OR ≥3 days/week participating in a resistance training program AND the ability to squat ≥ 1.5x body mass (males) or ≥ 1.2x body mass (females) (T group). Following verbal and written consent, participants then completed testing procedures described in greater detail in the following sections. Descriptive characteristics of participants are presented below in Table 1.

**Table 1.** Pre-intervention participant characteristics.

| Characteristic | Overall (*n* = 21) | Trained (*n* = 10) | Untrained (*n* = 11) |
| --- | --- | --- | --- |
| Height (cm) | 174 ± 7 | 177 ± 8 | 172 ± 5 |
| Sex | 16 M/ 5 F | 8 M/ 2 F | 8 M/ 3 F |
| Body Mass (kg) | 78.6 ± 18.3 | 81.4 ± 8.1 | 78.4 ± 24.4 |
| Age (years) | 26 ± 3 | 27 ± 3 | 26 ± 2 |
| Self-reported training (years) | --- | 6 ± 4 | --- |
| Body Fat % | 25.4 ± 8.9 | 20.8 ± 8.9 | 30.5 ± 6.9* |
| VL mCSA (cm<sup>2</sup>) | 28.1 ± 10.1 | 32.7 ± 9.4 | 25.1 ± 9.1* |
| VL Thickness (cm) | 2.5 ± 0.5 | 2.8 ± 0.3 | 2.4 ± 0.5* |
| 1RM leg press (kg) | 180 ± 126 | 270 ± 113 | 103 ± 70* |
| 1RM hexbar deadlift (kg) | 100 ± 46 | 135 ± 39 | 70 ± 23* |

Legend: Data are presented as mean ± standard deviation values. \*, indicates a significant pre-intervention difference based on an independent samples t-test.

### Familiarization/Squat Testing Session

All participants visited the laboratory to undergo a familiarization session. During this session, participants were exposed to the assessments to occur during subsequent data collection sessions. Participants in the UT group were then acclimated to each RT exercise that was to be performed during the post-disuse recovery phase. RT technique was taught by trained and two licensed certified strength and conditioning specialists who were members of the study team. Participants were deemed suitable for participation when they demonstrated proper technique. Participants in the T group were also shown to the weight room, where they completed a warmup of their choice and then demonstrated the ability to barbell back squat ≥1.5x their own body mass (males) or ≥1.2x their own body mass (females). T participants were scheduled for follow-up testing sessions after proving proficiency in the barbell back squat and lifting technique with all other involved exercises.

### Study Design Overview

The experimental design is presented in Figure 1. After the familiarization session, T and UT participants underwent 2 weeks of left leg immobilization via a locking leg brace and crutches followed by eight weeks of RT. Participants visited the laboratory for data collection sessions prior to commencement of the study (PRE), after the disuse protocol (MID) and after 8 weeks of RT (POST). At each data collection session, participants underwent measurements of height (PRE only), body mass, urine specific gravity, bioelectrical impedance spectroscopy (BIS) to assess body fat percentage, muscle thickness and CSA via ultrasound of the left vastus lateralis (VL), a biopsy of the left VL, and three repetition maximum tests (3RM) of the hex-bar deadlift and the leg press. The order of testing was the same for all three time points. PRE and POST occurred a minimum of 72 hours after any habitual RT bout in all participants, and POST also occurred ≥ 72 hours after the last study prescribed RT bout. MID occurred immediately after the removal of the locking leg brace. Participants were compensated $1200 for participation in this study.

**Figure 1.**
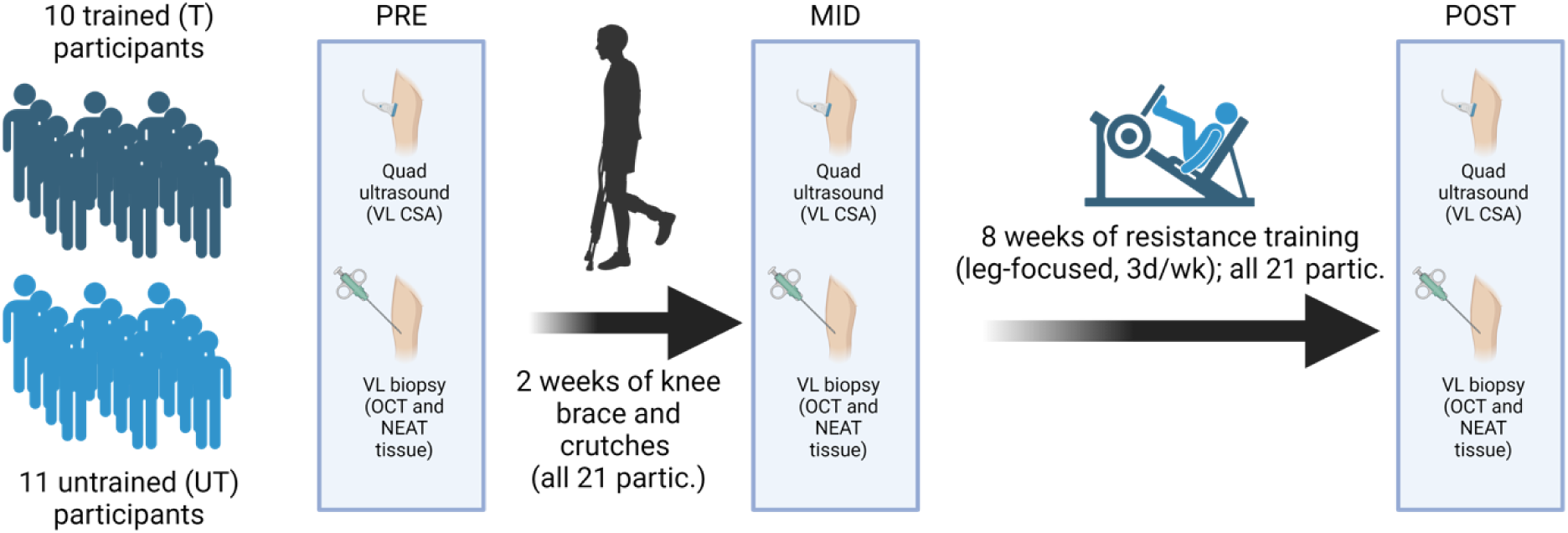
Study design. Legend: Summary of study design. Schematic (drawn using Biorender.com) illustrates the study logistics and participant number for this study. More details regarding study design can be found in the text. Abbreviations: VL, vastus lateralis; CSA, cross-sectional area (note, presented as VL mCSA in-text); OCT, optimal cutting temperature.

### Leg Disuse Protocol

The leg disuse protocol was designed based on previous work by MacLennan et al. (25). Upon completion of all testing procedures during PRE, participants were fitted for a knee joint immobilizer brace (T Scope® Premier Post-Op Knee Brace; Breg Inc., Carlsbad, CA, USA). The brace extended from midcalf to midthigh and was held in place with a series of padded Velcro straps. The brace was locked in place at ∼90° of knee flexion. In this position the participants’ knee extensors remained relaxed and unloaded to ensure the leg was fully non-weight bearing. Participants were additionally fitted for axillary crutches and were provided gait-training to effectively ambulate with a single leg while using crutches. All crutch-fitting and gait-training were carried out by a licensed physical therapist that was a part of the study team. Participants were also provided with a compression stocking to mitigate the risk of skin lesions (e.g. rashes) or potential clotting issues induced by two weeks of leg bracing and provided guidance on skincare and hygiene. Notably, no adverse events were reported by participants in this regard. Participants were instructed to always wear the brace except when in the shower or preparing to sleep for the night. After 14 days of bracing and ambulation on crutches, each participant returned to the laboratory for MID testing. At this time the brace was removed, and each participant returned to their bilateral full weight bearing gait.

### Resistance Training Protocol

Following removal of the leg brace and MID testing, participants began an eight-week RT block by performing RT three nonconsecutive days per week (typically Monday, Wednesday, and Friday). The exercises performed were as follows: Monday and Friday Session: Barbell back squat, leg extension machine, barbell bench press, lat pulldowns, lying hamstring curl; Wednesday Session: leg press, hex-bar deadlift, barbell row, barbell overhead press, dumbbell bicep curl. Details regarding the progression model of this intervention are presented in Table 2. Although the prescription priority was to target the knee extensors, exercises for most all major muscle were additionally included to improve adherence and decrease likelihood of participants performing RT outside of study protocols. Only those exercises targeting the knee extensors (barbell back squat, hex-bar deadlift, leg press, and leg extension machine) progressed in volume throughout the study. Set number was progressed throughout the duration of the study for those exercises targeting the knee extensors. Hex-bar deadlift and leg press loads were prescribed based on predicted 1RMs from the 3RM tests while the loads for the remainder of the exercises were prescribed based on rating of perceived exertion (RPE). For those exercises performed based on RPE, participants were instructed to perform each exercise at an approximate RPE of 8 out of a maximum 10 value. This is based on the OMNI RPE scale where a value of 8 corresponds to a hard lift (26).

**Table 2.** Progression model for resistance training intervention.

| Week | % estimated 1RM/RPE | Repetition Range | Number of Sets* |
| --- | --- | --- | --- |
| 1 | 65/8 | 8-12 | 3** |
| 2 | 70/8 | 8-12 | 3 |
| 3 | 65/8 | 8-12 | 4 |
| 4 | 65/8 | 8-12 | 4 |
| 5 | 65/8 | 8-12 | 5 |
| 6 | 65/8 | 8-12 | 6 |
| 7 | 65/8 | 8-12 | 7 |
| 8 | 65/8 | 8-12 | 8 |

Abbreviations: % 1RM, Percentage of pre-tested one repetition maximum for a given exercise; \*Total number of sets for exercises targeting the knee extensors; \*\*Exercises not targeting the knee extensors were held constant at 3 sets per exercise.

**Table 3.** Antibodies used for western blotting.

| Target | Host Species | Catalog Number | Manufacturer |
| --- | --- | --- | --- |
| Calpain 1 | Rabbit | 2556 | Cell Signaling Technology |
| Calpain 2 | Rabbit | 70655 | Cell Signaling Technology |
| LC3 | Rabbit | 2775 | Cell Signaling Technology |
| p62 (SQSTM1) | Rabbit | 5114 | Cell Signaling Technology |
| Beclin-1 | Rabbit | NB110-8731855 | Novus Biologicals |
| Phospho-ULK (Ser555) | Rabbit | 5869 | Cell Signaling Technology |
| ULK | Rabbit | 8054 | Cell Signaling Technology |
| Ubiquitin | Rabbit | 3933 | Cell Signaling Technology |
| cMyc | Rabbit | 9402 | Cell Signaling Technology |
| SKIV2L2 | Mouse | Sc-515828 | Santa Cruz Technology |
| ExoSC10 | Mouse | Sc-374595 | Santa Cruz Technology |
| G3BP1 | Mouse | Sc-365338 | Santa Cruz Technology |
| USP10 | Rabbit | 5553 | Cell Signaling Technology |
| UBF | Rabbit | Ab109011 | Abcam |
| POLR1B | Rabbit | PA5-88309 | Thermo Fisher |
| Phospho-IRE1α (Ser51) | Rabbit | PA1-16927 | Thermo Fisher |
| IRE1α | Rabbit | 3294 | Cell Signaling Technology |
| ATF4 | Goat | Ab1371 | Abcam |
| Xbp1s | Rabbit | 40435 | Cell Signaling Technology |
| ATF6 | Mouse | ALX-804-381 | Enzo Life Sciences |
| CHOP | Mouse | 2895 | Cell Signaling Technology |
| BiP (GRP78) | Rabbit | 3183 | Cell Signaling Technology |
| Phospho-eIF2α (Ser51) | Rabbit | 3398 | Cell Signaling Technology |
| eIF2α | Rabbit | 5324 | Cell Signaling Technology |
| Phospho-P70S6K1 (Thr389) | Rabbit | 2983 | Cell Signaling Technology |
| P70S6K1 | Rabbit | 9202 | Cell Signaling Technology |
| Phospho-RPS6 (Ser235/236) | Rabbit | 4858 | Cell Signaling Technology |
| RPS6 | Rabbit | 2217 | Cell Signaling Technology |
| Phospho-4EBP1 (Thr37/46) | Rabbit | 2855 | Cell Signaling Technology |
| 4EBP1 | Rabbit | 9644 | Cell Signaling Technology |

Abbreviations: LC3, microtubule-associated protein 1B-light chain 3; p62, sequestosome 1; ULK, Unc-51-like kinase 1; cMyc, myc proto-oncogene; SKIV2L2, superkiller viralicidic activity 2-like 2; ExoSC10, exosome component 10; G3BP1 stress granule assembly factor 1; USP10, ubiquitin specific peptidase 10; UBF, upstream binding factor; POLR1B, RNA polymerase 1 subunit B; IRE1α , inositol requiring enzyme 1 alpha; ATF4, activating transcription factor 4; Xbp1s, spliced form of X-box binding protein 1; ATF6, activating transcription factor 6; CHOP, CCAAT enhancer binding protein (C/EBP) homologous protein; BiP, binding immunoglobulin protein; eIF2α , eukaryotic initiation factor 2 alpha; P70S6K1, ribosomal protein S6 kinase beta-1; RPS6, ribosomal protein S6; 4EBP1, eukaryotic translation initiation factor 4E-binding protein 1; phospho-, phosphorylated.

### Testing Sessions (PRE, MID, and POST)

#### Urine Specific Gravity, Height, and Body Mass

During testing session visits, participants reported to the laboratory following an overnight fast. Hydration status was assessed via a urine specific gravity test measured with a handheld refractometer (ATAGO; Bellevue, WA, USA) with the goal of ensuring adequate hydration (value ≤1.020) (27). Body mass and height were assessed with a digital scale (Seca 769, Hanover, MD, USA).

#### Ultrasonography

Ultrasonography was then performed to determine the thickness and cross-sectional area of the left VL as described previously by our laboratory (Ruple, Mesquita et al. 2022). All measurements were taken at the midway point between the iliac crest and the proximal patella by the same member of the research team.

#### Vastus Lateralis Muscle Biopsies

After ultrasound scans, skeletal muscle biopsy samples were collected from the left VL at the same location as ultrasound imaging using a 5 mm Bergstrom needle. Biopsies at MID and POST were taken ∼2 cm proximal from the preceding biopsy scar. Briefly, participants laid supine on an athletic training table and the upper thigh was shaved and cleaned with 70% isopropanol prior to receiving an injection of 1% lidocaine (0.6-0.9 mL). After waiting ∼5 minutes to allow lidocaine to take full effect, the area was cleaned with chlorhexidine and a pilot incision made with a sterile, single-use No. 11 surgical blade (AD Surgical, Sunnyvale, CA, USA). The biopsy was then collected using a 5 mm Bergstrom needle under suction. Approximately 50-100 mg of skeletal muscle tissue was collected, immediately teased of blood and connective tissue, and separated for histological and biochemical analysis. Tissue (∼30 mg) was mounted for histology at a 90° angle on a piece of cork in a 1:1 w/w mixture of optimal cutting temperature (OCT) solution and tragacanth powder (Alfa Aesar, Ward Hill, MA, USA). The mount was then covered in OCT, frozen in liquid nitrogen cooled 2-methylbutane for ∼30 seconds, then contained in a box top floating atop liquid nitrogen before long-term storage at -80°C. A separate ∼20-60 mg tissue sample was placed in foil and flash frozen in liquid nitrogen for downstream biochemical analyses. All tissue triage procedures occurred within a 2-minute window.

#### Strength Testing

Following biopsies, participants were escorted to the weight room to perform a standardized warm-up and maximal strength testing. Briefly, two research team members conducted all strength testing and were present for all testing. Participants were prescribed a warm-up based on RPE (PRE) or previous weights lifted (MID, POST) wherein ∼3-5 warm up sets were achieved ranging from 4-8 on the RPE scale (PRE) or 40%-80% 3RM (MID and POST) prior to the first attempt at a 3RM. Following warm-ups, participants were allowed three attempts to attain the highest 3RM possible with technique being maintained in the leg press and the hex-bar deadlift. Weights achieved were then transformed via the National Strength and Conditioning Association RM conversion calculator for conversion to a predicted 1RM for statistical analysis.

### Muscle tissue processing and analytical techniques

#### Western Blotting

Muscle tissue was lysed using a general cell lysis buffer (Cell Signaling Technology, Danvers, MA, USA; Cat. No. 9803) and tight-fitting pestles. Subsequent lysates were processed for total protein using a commercially available BCA protein assay kit (Thermo Fisher, Waltham, MA, USA; Cat. No. A55864) and spectrophotometer (Agilent Biotek Synergy H1 hybrid reader; Agilent, Santa Clara, CA, USA). Lysates were then centrifuged at 500 g for 5 minutes and supernatants were prepared for western blotting using 4x Laemmli buffer and deionized water (diH2O) at equal protein concentrations (1 μg/μL). 15 μL of samples were pipetted onto SDS gels (4-15% Criterion TGX Stain-free gels, Bio-Rad Laboratories; Hercules, CA, USA), and proteins were separated by electrophoresis at 200 V for 45-50 min. Proteins were then transferred to methanol-preactivated PVDF membranes (Bio-Rad Laboratories) for 2 h at 200 mA, Ponceau stained for 10 minutes, washed with diH2O for ∼30 seconds, dried, and digitally imaged (ChemiDoc Touch, Bio-Rad). Following Ponceau imaging, membranes were reactivated in methanol, blocked with non-fat bovine milk for ≥1 hour, and washed 3x5 minutes in tris-buffered saline with tween 20 (TBST). Membranes were incubated with primary antibodies (1:1000 v/v/ dilution in TBST with 5% bovine serum albumin (BSA)) on a rocker overnight, or for 48 hours (phosphorylated targets) at 4° C. Primary antibodies used to detect protein targets are listed below in Table 4.

Following primary antibody incubations, membranes were washed 3x5 min in TBST and incubated for 1 h with HRP-conjugated anti-rabbit IgG (Cell Signaling Technology, Cat No. 7074), HRP-conjugated anti-mouse IgG (Cell Signaling Technology, Cat No. 7076), or HRP-conjugated anti-goat IgG (Genetex, Irvine, CA, USA Cat No. GTX628547-01). Membranes received a final set of 3x5 min washes in TBST, then developed using chemiluminescent substrate (Millipore, Burlington, MA, USA), and then digitally imaged. Raw target band densities were obtained and normalized to Ponceau densitometry values. For non-phosphorylated targets, fold-change values were derived by dividing Ponceau-normalized band density values by the aggregate PRE mean value of the UT group for each target. Band density values for phosphorylated targets were divided by corresponding pan band density values, and again fold-change values were derived by dividing these values by the aggregate PRE mean value of the UT group.

#### Calpain and 20S Proteasome Activity

Calpain activity was measured via commercially available kit (Promega, Madison, WI, USA; Cat No. G8502) according to manufacturer’s instructions. Briefly, 50 μL of sample diluted at 1:10 in diH2O was combined with 50 μL of Calpain-Glo™ reagent and CaCl2 (2 mM) in a 96-well white plate to produce a final reaction volume of 100 μL. The assay blank was included consisting of 100 μL of Calpain-Glo™ reagent and 2 mM CaCl2 without a protein sample. The plates were then orbitally shaken at 400 RPM for 30s, incubated at room temperature for 15 minutes and read with a luminometer with the following settings: Gain=135, Read height=1 mm, Exposure=20s. Relative expression units (REUs) were generated, and the value of the blank was subtracted from each sample to control for background. These values were then normalized to total protein loaded and analyzed.

Chymotrypsin-like 20S Proteasome activity was measured via a commercially available kit (Promega, Madison, WI, USA; Cat No. G8622) according to manufacturer’s instructions. Briefly, 50 μL of samples diluted at 1:10 with diH2O were combined with 50 μL of Proteasome-Glo™ reagent in a 96-well white plate to produce a final reaction volume of 100 μL. The assay blank was 100 μL of Proteasome-Glo™ reagent without a protein sample. The plates were then orbitally shaken at 400 RPM for 30s, incubated at room temperature for 15 min and then read with a luminometer with the following settings: Gain=135, Read heigh=1 mm, Exposure=20 s. REUs were then generated, and the value of the blank was subtracted from each sample to control for background. These values were then normalized to total protein loaded and analyzed.

#### RNA Isolation and Quantification of Total RNA, 18S rRNA and 28S rRNA

RNA was isolated via a commercially available kit (Qiagen, Hilden, Germany; Cat. No. 74004) using Qiazol (Qiagen, Cat. No. 79306) in accordance with manufacturer instructions. Briefly, approximately 20 mg of tissue were placed in 500 μL of Qiazol and homogenized with a bead homogenizer (Bead Ruptor ELITE; OMNI International, Kennesaw, GA, USA). Samples were then passed through column filtration as recommended by manufacturer’s protocol. Final RNA products were then eluted in 20 μL of DEPC-treated H2O and subject to further analysis.

Total RNA concentration was determined via a commercially available assay (Thermo Fisher Scientific, Cat. No. Q33211) and measured using a Qubit 4.0 Analyzer (Thermo Fisher Scientific). 18S and 28S rRNAs were determined with a commercially available assay kit (Agilent, Santa Clara, CA, USA; Cat. No: 5067-5576) and analyzed using an Agilent 4200 TapeStation per manufacturer’s instructions.

#### Immunohistochemistry for fCSA Determination

OCT preserved samples were sectioned at a thickness of 12 μm using a cryostat (Leica Biosystems, Buffalo Grove, IL, USA), adhered to positively charged histology slides, and stored at -80° C until batch-processed for immunohistochemical analyses to measure type I and type II fiber CSA (fCSA). All samples from the same participant were placed on the same slide and analyzed concomitantly.

Slides were stained for the visualization of dystrophin and type I myosin heavy chain. The slides were removed from -80° C storage and frozen cryosections were air-dried at room temperature for ≥2 h and fixed with acetone at -20 °C for 5 minutes. Slides were then incubated with 3% H2O2 for 10 min at room temperature, followed by a 1-minute incubation with autofluorescence quenching reagent (TrueBlack, Cat. No. 23007; Biotium, Fremont, CA, USA) and blocked for 1 hour with a 5% goat serum, 2.5% horse serum, and 0.1% Triton-X solution at room temperature. After blocking, slides were incubated overnight at 4°C with a primary antibody cocktail containing 1:100 dystrophin (Cat. No. GTX57970; GeneTex) + 1:100 BA-D5 (myosin heavy chain I, Cat. No. BA-D5; Developmental Studies Hybridoma Bank, Iowa City, IA, USA) + 2.5% horse serum in phosphate-buffered saline (PBS). The following day, the primary antibody was removed, slides were rinsed with PBS, and incubated for 60 minutes in a secondary antibody cocktail containing 1:250 goat anti-mouse IgG2b Alexa Fluor 647 (Cat. No. A-21242; Thermo Fisher Scientific) + 1:250 goat anti-rabbit IgG DyLight488 (cat. no. DI-1488; Vector Laboratories, Newark, CA, USA) in PBS. Slides were then stained with DAPI (1:10,000; 4’ ,6-diamidino-2-phenylindole; cat. no. D3571; Thermo Fisher Scientific) for 10 minutes at room temperature and mounted with glass coverslips using 1:1 PBS and glycerol as mounting medium. Sections were stored in the dark at 4 °C until imaging was conducted. Digital images were captured with a fluorescence microscope at (Zeiss Axio Imager.M2) and motorized stage. All areas selected for analyses were free of freeze-fracture artifacts, and an average of 466 fibers were considered for analysis. Open-sourced software (Myovision) was used to analyze all images for mean fCSA, fiber type, fiber type specific fCSA, and nuclei per fiber (28). All images were manually checked after going through Myovision’s analysis and two images (one T MID, one UT POST) were removed due to poor section quality and the inability for the software to correctly delineate fiber borders in these samples.

#### Satellite Cell Number Determination

Separate slides were stained for the visualization of satellite cells in accordance with previous published methods from our laboratory (29). Briefly, sections were fixed in ice-cold acetone for 5 minutes, incubated in 3% hydrogen peroxide for 10 minutes and 1x True Black for 1 minute to block autofluorescence. Sections were then blocked for 1 hour with 5% goat serum/2.5% horse serum/0.1% Triton-X 100, followed by streptavidin solution and biotin solution blocks for 15 minutes each. Sections were incubated overnight at 4°C with the following primary antibody cocktail: mouse anti-Pax-7 IgG1 (1:20) (supernatant; Developmental Studies Hybridoma Bank; Iowa City, IA), rabbit anti-dystrophin (1:100; GTX57970; GeneTex), and 2.5% horse serum. The following day the primary antibody cocktail was removed, slides were rinsed in PBS, and were incubated with goat anti-mouse IgG1 biotin-SP-conjugated (1:1,000) (Jackson ImmunoResearch; West Grove, PA, USA) in 2.5% horse serum for 90 minutes. Sections were then incubated in a secondary antibody cocktail that included SA-HRP (1:500) (Thermo Fisher Scientific; Cat. No. S-911) and goat anti-rabbit AF488 (1:250) (Vector Laboratories; Burlingame, CA) for 1 hour. TSA AF594 (1:200) (Thermo Fisher Scientific; Cat. No.: B-40957) was added for 20 minutes, followed by DAPI (1:10,000) for 10 minutes. Coverslips were added using PBS + glycerol as mounting medium, and entire sections at ×20 objective digital images were collected for each participant sampling time point were captured with a fluorescent microscope (Zeiss Axio Imager.M2). Satellite cell analysis was performed manually by a single investigator who used a hand tally counter to quantify cells outside the dystrophin border co-stained with Pax7 and DAPI (Pax7+/DAPI+). Data are presented as satellite cells per 100 fibers, and all fibers present in a given section were analyzed per sample.

### Statistical Analyses

Data were plotted and analyzed in GraphPad Prism (v10.2.2). All data were checked for normality via Shapiro-Wilk tests and analyzed via 2 x 3 (group*time) two-way ANOVAs or mixed-effect models. Tukey’s *post hoc* tests were used to determine pairwise differences between group and/or timepoints if interaction term was significant. Simple main effects analyses were performed if only a main effect was significant. Notably, total volume load data were analyzed for normality and analyzed via unpaired student’s t-test. Statistical significance was established as P≤0.05, and all data are expressed as mean and standard deviation with individual data points shown throughout.

## RESULTS

### Phenotype and Performance Outcomes

Hex-bar deadlift predicted 1RM values demonstrated main effects of time and training status (P≤0.004) and a significant training status*time interaction (P=0.035; Figure 2a). T had greater hex-bar deadlift predicted 1RM values than UT at all time points (P≤0.002) and both groups did not significantly decrease from PRE to MID (P≥0.063). Both groups did, however, increase at POST as compared to both PRE (P≤0.004) and MID (P≤0.001). Leg press predicted 1RM values demonstrated significant main effects of time and training status (P≤0.002) and a significant interaction (P=0.044; Figure 2b). T had greater leg press predicted 1RM values at all time points than UT (P≤0.023). T decreased leg press predicted 1 RM from PRE to MID (P=0.013) and increased at POST from both PRE and MID (P≤0.022). UT however did not decrease significantly from PRE to MID (P=0.995) but increased at POST as compared to both PRE and MID (P<0.001). Ultrasound-derived VL thickness demonstrated a main effect of time and a training status*time interaction (P≤0.015) without a main effect of training status (P=0.061; Figure 2c). VL thickness was greater at PRE in T than UT (P=0.011). Both T and UT demonstrated significant declines from PRE to MID (P≤0.038), both groups increased from MID to POST (P<0.001), but only UT demonstrated increases from PRE to POST (P<0.001). VL mCSA demonstrated a significant main effect of time and a significant time*training status interaction (P≤0.014), but no main effect of training status (P=0.069; Figure 2d). T demonstrated greater VL mCSA at PRE and MID than UT (P≤0042). T and UT showed significant decreases in VL mCSA from PRE to MID (P≤0.044) followed by significant increases from MID to POST (P≤0.002). Only UT demonstrated a significant increase from PRE to POST (P<0.001). Total volume load (reps*sets*load) for the duration of the study was greater in T versus UT (P<0.001; Figure 2e).

**Figure 2.**
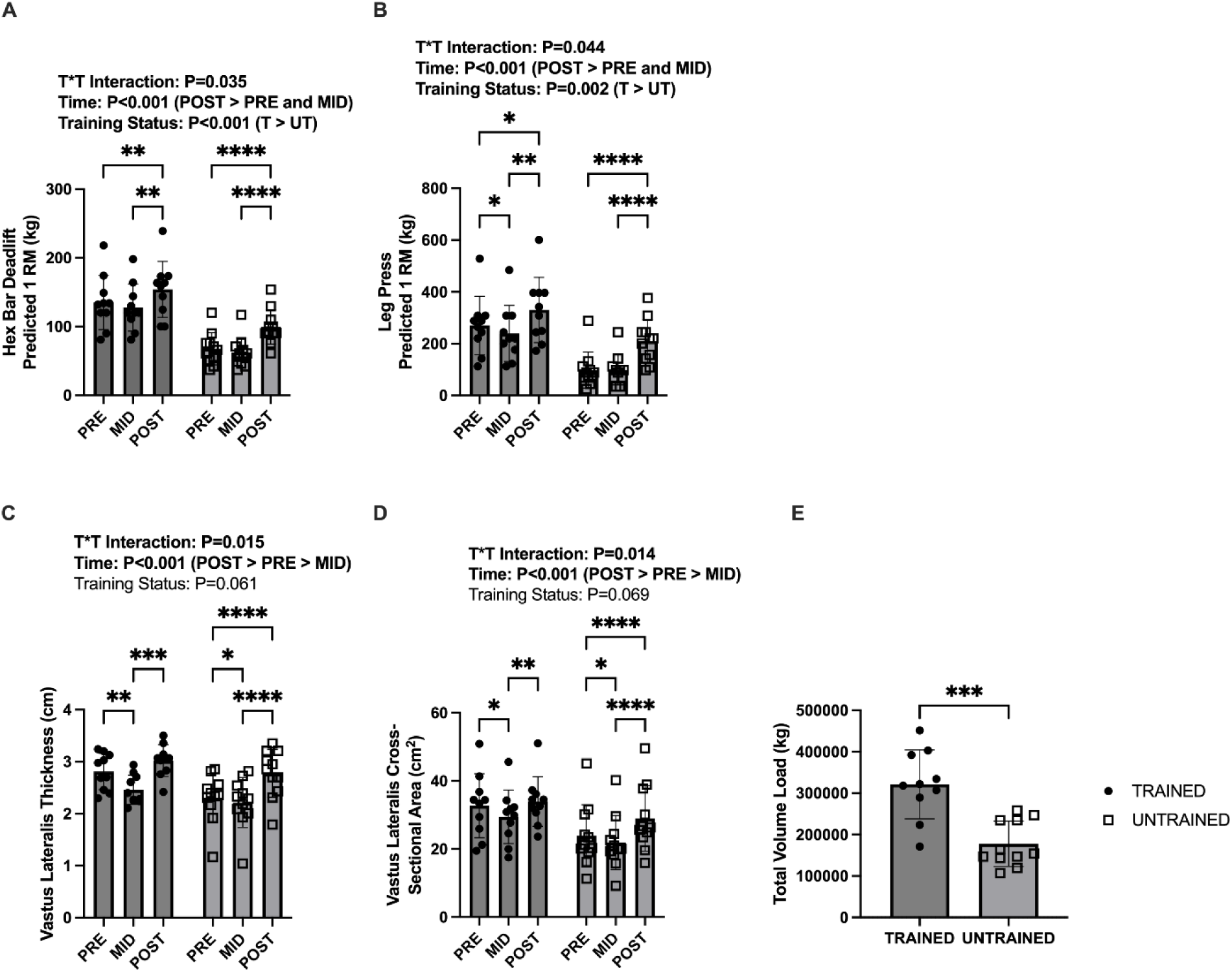
VL morphology and strength responses. Legend: Data are presented as individual data points with mean and standard deviation bar graphs. Results from 21 human participants for hex bar deadlift predicted 1 repetition maximum (panel a), leg press predicted 1 repetition maximum (panel b), ultrasound-derived vastus lateralis thickness (panel c), ultrasound-derived vastus lateralis cross-sectional area (panel d), and total volume load throughout 8 weeks of resistance training (panel e) are presented. Two-way ANOVA-derived P values for training status*time interactions, main effects of time, and main effects of training status are presented above graphs. For significant main effects, *post hoc* test details are presented in parentheses (panels a-d). *Post hoc* statistical significance between time points is presented for those ANOVAs demonstrating a significant training status*time interaction (panels a-d). T-test statistical significance is demonstrated in panel e. Symbols: *, P<0.05; **, P<0.01; ***, P<0.005; ****, P<0.001. Abbreviations: 1 RM, 1 repetition maximum.

### Fiber Cross-Sectional Area and Fiber Type Data

Mean fCSA did not demonstrate a training status*time interaction (P=0.279; Figure 3a), but did demonstrate a main effect of time (P=0.002) where PRE and POST were greater than MID, and a main effect of training status (P=0.012) where T was greater than UT. Type I fiber percentage did not demonstrate main effects of training status or time (P≥0.476) nor a training status*time interaction (P=0.114, Figure 3b). Type I fCSA did not demonstrate main effects of time or training status (P≥0.142) nor a training status*time interaction (P=0.252, Figure 3c). Type II fCSA did not demonstrate a training status*time interaction (P=0.370; Figure 3d), but did demonstrate a main effect of time (P<0.001) where PRE and POST were greater than MID, and training status (P=0.003) where T was greater than UT.

**Figure 3.**
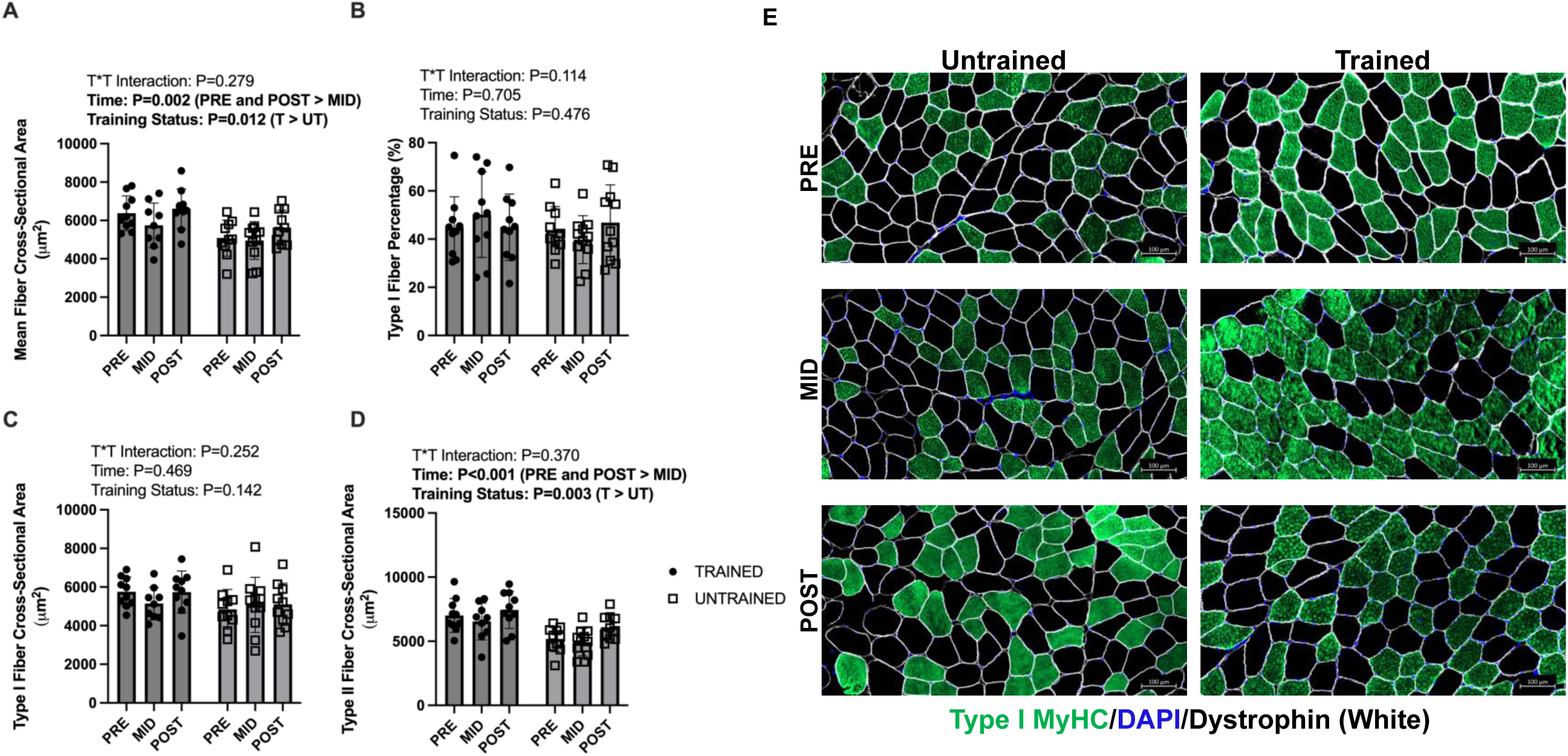
Skeletal muscle fiber level responses. Legend: Data are presented as individual data points with mean and standard deviation bar graphs. Results from 21 human participants for mean fiber cross-sectional area (panel a), type I fiber percentage (panel b), type I muscle fiber cross-sectional area (panel c), and type II muscle fiber cross-sectional area (panel d) are presented. Representative images for immunohistochemical staining are presented in panel e. Two-way ANOVA-derived P values for training status*time interactions, main effects of time, and main effects of training status are presented above graphs. For significant main effects, *post hoc* test details are presented in parentheses. Abbreviations: MyHC, Myosin Heavy Chain; DAPI, 4’,6-diamidino-2-phenylindole.

### Nuclear and Satellite Cell Data

Total nuclei per fiber did not demonstrate a main effect of time or training status*time interaction (P≥0.602; Figure 4a) but did exhibit a main effect of training status (P=0.004) where T was greater than UT. Type I fiber myonuclear number did not demonstrate a main effect of time or training status (P≥0.089) nor a training status*time interaction (P=0.404, Figure 3g). Type II fiber myonuclear number did not demonstrate a main effect of time nor a training status*time interaction (P≥0.562, Figure 3h), but did exhibit a main effect of training status (P=0.001) where T was greater than UT. Satellite cells per 100 fibers did not demonstrate a training status*time interaction (P=0.896; Figure 4b), but did demonstrate main effects of time (P<0.001) where POST was greater than PRE and MID, and training status where T was greater than UT.

**Figure 4.**
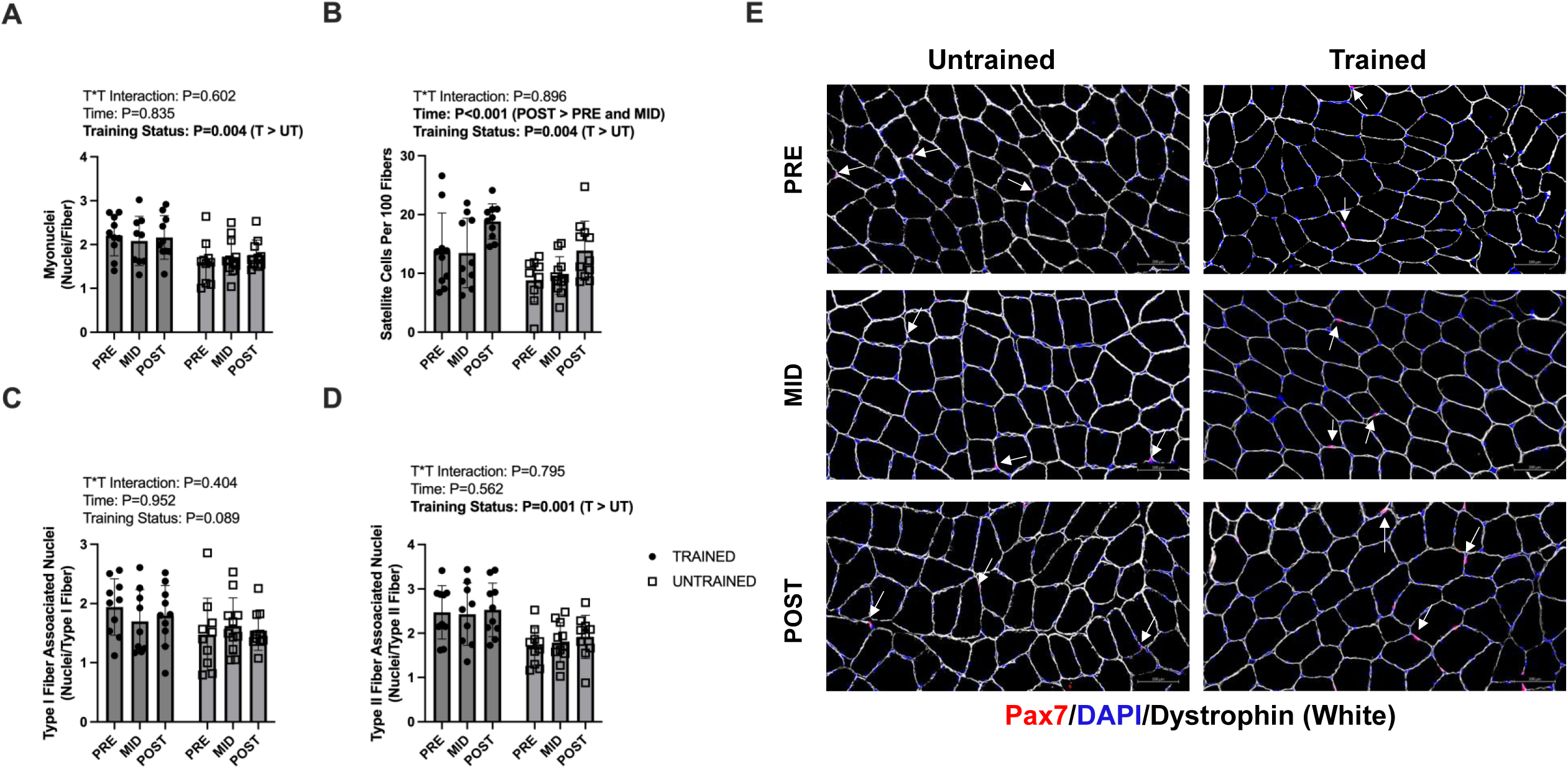
Myonuclear number and satellite cell responses. Legend: Data are presented as individual data points with mean and standard deviation bar graphs. Results from 21 human participants for total myonuclei (panel a), satellite cells per 100 fibers (panel b), Type I muscle fiber associated nuclei (panel c), and Type II muscle fiber associated nuclei (panel d) are presented. Representative images for immunohistochemical staining with arrows denoting Pax7+ cells are presented in panel e. Two-way ANOVA-derived P values for training status*time interactions, main effects of time, and main effects of training status are presented above graphs. For significant main effects, *post hoc* test details are presented in parentheses. *Post hoc* statistical significance between time points is presented for those ANOVAs demonstrating a significant training status*time interaction (panels a-d). Symbols: *, P<0.05; **, P<0.01; ***, P<0.005; ****, P<0.001. Abbreviations: Pax7, Paired Box 7; DAPI, 4’,6-diamidino-2-phenylindole.

### Proteolytic Markers

Calpain-1 protein expression exhibited a main effect of time (P=0.001) where PRE and POST were greater than MID but did not demonstrate a main effect of training status or training status*time interaction (P≥0.845; Figure 5a). Calpain-2 protein expression did not demonstrate a training status*time interaction (P=0.359; Figure 5b), but did demonstrate main effects of time (P<0.001) where POST > PRE > MID, and training status (P=0.029), where T was greater than UT. Calpain activity did not demonstrate a main effect of time or training status (P≥0.062) and also did not demonstrate a training status*time interaction (P=0.581; Figure 5c). p62 (Figure 5e) and poly-ubiquitinated protein expression (Figure 5h) exhibited a main effect of time (P≤0.029) where poly-ubiquitinated proteins were greater at POST than PRE. Neither p62 nor poly-ubiquitinated proteins demonstrated main effects of training status or a training status*time interaction (P≥0.097). LC3 II/I (Figure 5d), Beclin-1 (Figure 5f), and phosphor/pan ULK did not exhibit main effects of time, training status, or a training status*time interaction (P≥0.058). 20S proteasome activity did not demonstrate main effects of training status or time (P≥0.313) nor a training status*time interaction (P=0.087; Figure 5i).

**Figure 5.**
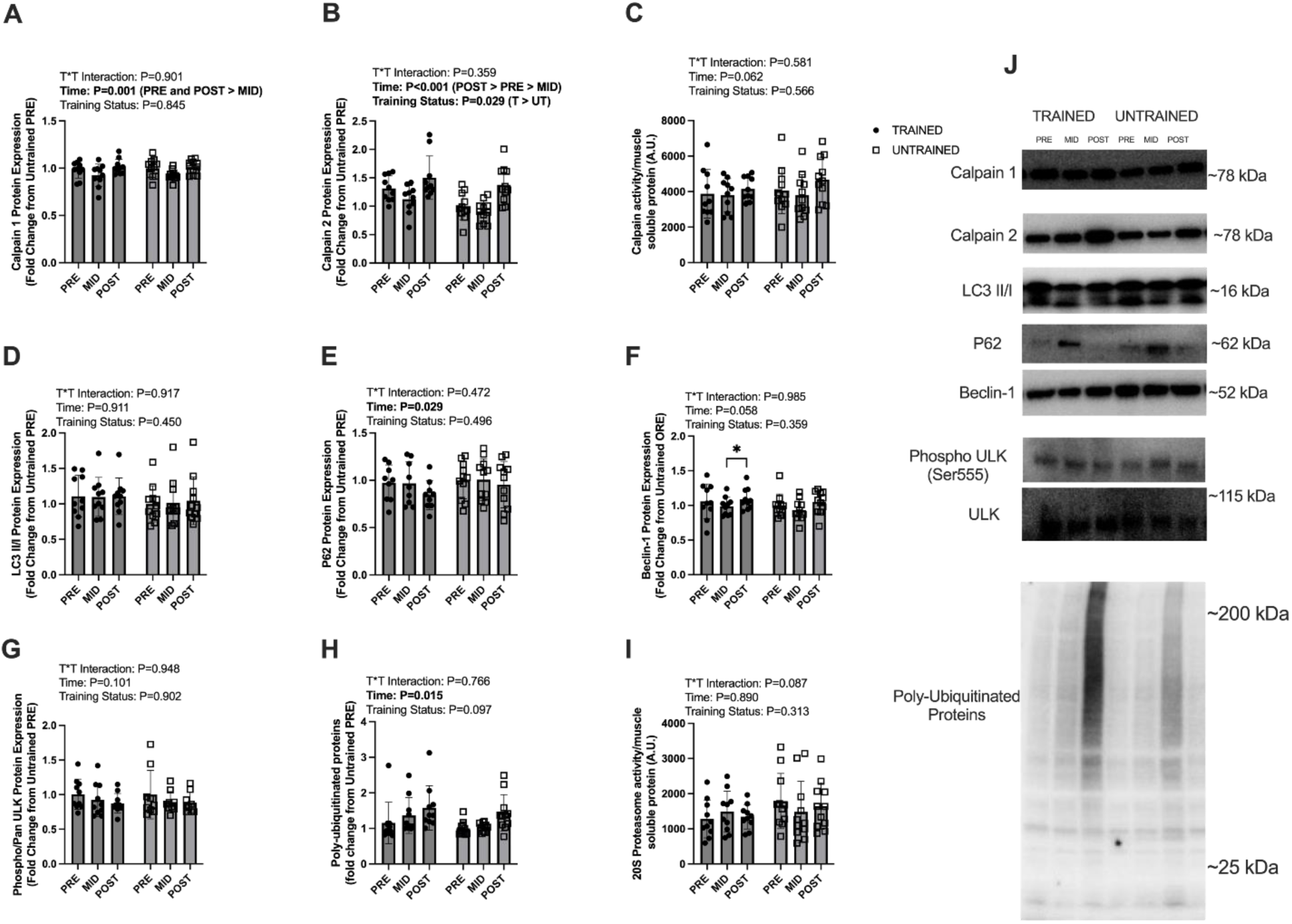
Proteolytic marker responses. Legend: Data are presented as individual data points with mean and standard deviation bar graphs. Results from 21 human participants for calpain-1 protein expression (panel a), calpain-2 protein expression (panel b), global calpain activity (panel c), LC3 II/I ratio expression (panel d), p62 protein expression (panel e), Beclin-1 protein expression (panel f), phosphorylated/pan ULK protein expression (panel g), poly-ubiquitinated protein levels (panel h), and 20S proteasome activity (panel i) are presented. Panel j contains representative western blot images. Two-way ANOVA-derived P values for training status*time interactions, main effects of time, and main effects of training status are presented above graphs. For significant main effects, *post hoc* test details are presented in parentheses. Abbreviations: LC3, microtubule-associated protein 1B-light chain 3; p62, sequestosome 1; ULK, Unc-51-like kinase 1; phospho-, phosphorylated.

### Ribosome Related Markers

Total RNA demonstrated a main effect of time (P<0.001), where PRE and POST were greater than MID, but no main effect of training status nor a training status*time interaction (P≥0.061; Figure 6a). 28S rRNA exhibited a main effect of time (P=0.008) where POST was greater than PRE and MID but did not exhibit a main effect of training status nor a training status*time interaction (P≥0.453; Figure 6b). Similarly, 18S rRNA demonstrated a main effect of time (P=0.007) where POST was greater than PRE and MID, without a main effect of training status nor training status*time interaction (P≥0.321; Figure 6c) cMyc protein expression did not demonstrate main effects of training status or time (P≥0.176) nor a training status*time interaction (P=0.271; Figure 6d). SKIV2L2 protein expression demonstrated a main effect of time (P=0.002) where PRE and MID were greater than POST but did not demonstrate a main effect of training status nor group*time interaction (P≥0.092; Figure 6e). Similarly, ExoSC10 protein expression demonstrated a main effect of time (P=0.002) where PRE and MID were greater than POST but did not demonstrate a main effect of training status nor training status*time interaction (P≥0.192; Figure 6f). G3BP1 protein expression also exhibited a main effect of time (P=0.017) where POST was greater than MID but did not demonstrate a main effect of training status nor a training status*time interaction (P≥0.082; Figure 6g). USP10 protein expression did not demonstrate main effects of time or training status (P≥0.061) nor a training status*time interaction (P=0.084; Figure 6h). UBF protein expression demonstrated a main effect of time (P<0.001) where PRE and MID were greater than POST but did not demonstrate a main effect of training status nor a training status*time interaction (P≥0.116; Figure 6i). POLR1B protein expression demonstrated a main effect of time (P=0.012) where MID was greater than PRE and POST but did not demonstrate a main effect of training status nor a training status*time interaction (P≥0.652; Figure 6j).

**Figure 6.**
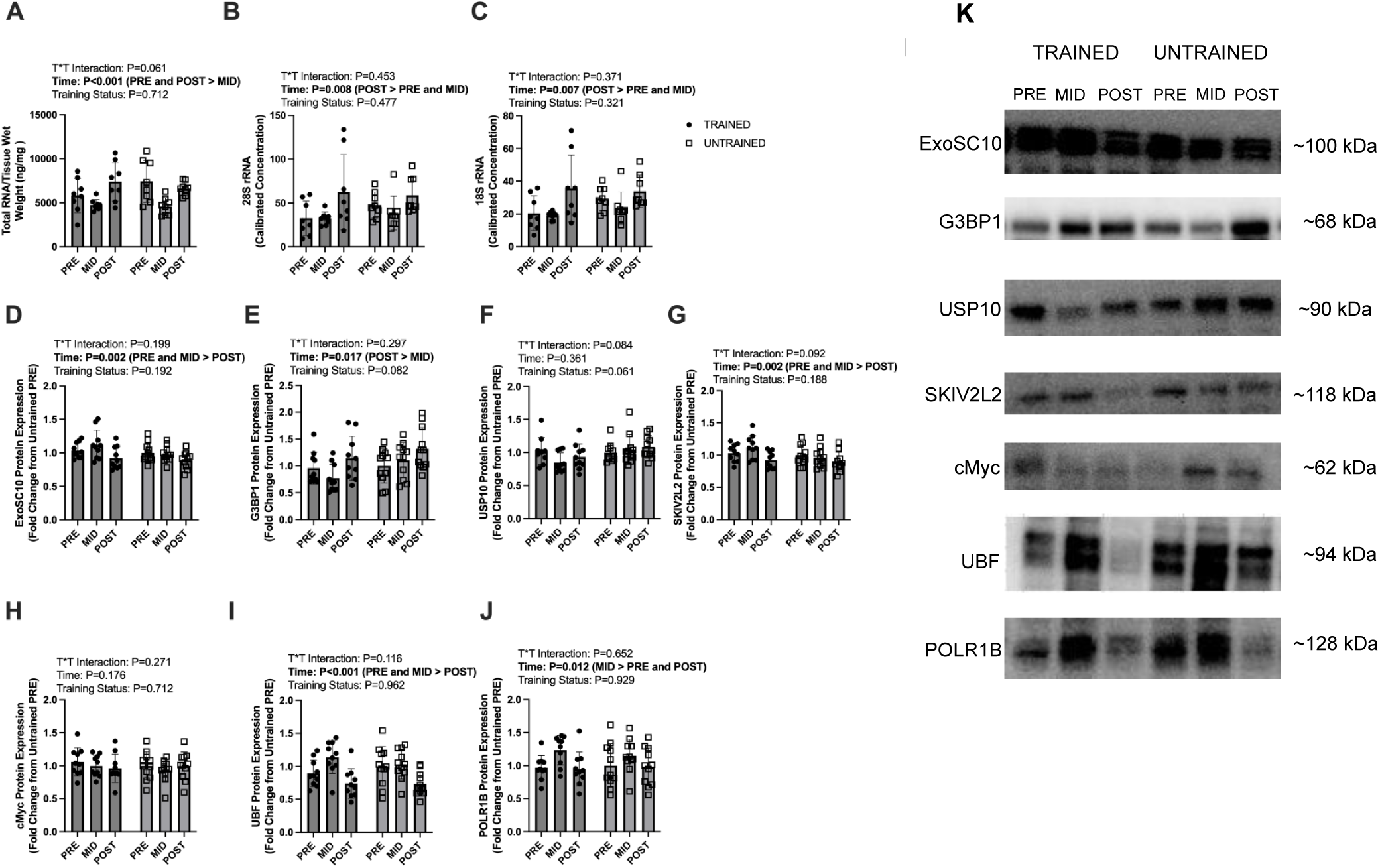
Ribosome content as well as ribosome biogenesis ribophagy marker responses. Legend: Data are presented as individual data points with mean and standard deviation bar graphs. Results from 21 human participants for total RNA (panel a), 28S rRNA concentration (panel b), 18S rRNA concentration (panel c), ExoSC10 protein expression (panel d), G3BP1 protein expression (panel e), USP10 protein expression (panel f), SKIV2L2 protein expression (panel g), cMyc protein expression (panel h), UBF protein expression (panel i), and POLR1B protein expression are presented. Panel k contains representative western blot images. Two-way ANOVA-derived P values for training status*time interactions, main effects of time, and main effects of training status are presented above graphs. For significant main effects, *post hoc* test details are presented in parentheses. Abbreviations: ExoSC10, exosome component 10; G3BP1 stress granule assembly factor 1; USP10, ubiquitin specific peptidase 10; SKIV2L2, superkiller viralicidic activity 2-like 2; cMyc, myc proto-oncogene; UBF, upstream binding factor; POLR1B, RNA polymerase 1 subunit B.

### Endoplasmic Reticulum Stress Markers

BiP protein expression demonstrated a main effect of time (P=0.002) where POST was greater than PRE and MID but did not demonstrate main effect of training status nor a training status*time interaction (P≥0.290; Figure 7a). Phospho/Pan IRE1α demonstrated a training status*time interaction (P=0.042), without main effects of group or time (P≥0.187; Figure 7b). Only UT demonstrated decrements at POST as compared to PRE and MID (P≤0.12). Xbp1s protein expression demonstrated a main effect of time (P=0.008) where POST was greater than MID but did not demonstrate a main effect of training status nor a training status*time interaction (P≥0.055; Figure 7c). Cleaved ATF6 did not demonstrate main effects of training status or time (P≥0.548) nor a training status*time interaction (P=0.905; Figure 7d). CHOP protein expression demonstrated main effects of time (P<0.001) where POST was greater than PRE and MID, and training status (P=0.038) where UT was greater than T, with no training status*time interaction (P=0.469; Figure 7e). ATF4 protein expression demonstrated a main effect of training status (P=0.048) where T was greater than UT without a main effect of time nor a training status*time interaction (P≥0.170; Figure 7f). Phospho/Pan eIF2α demonstrated a main effect of time (P=0.041) where MID was greater than POST but did not demonstrate a main effect of training status nor a training status*time interaction (P≥0.247; Figure 7g).

**Figure 7.**
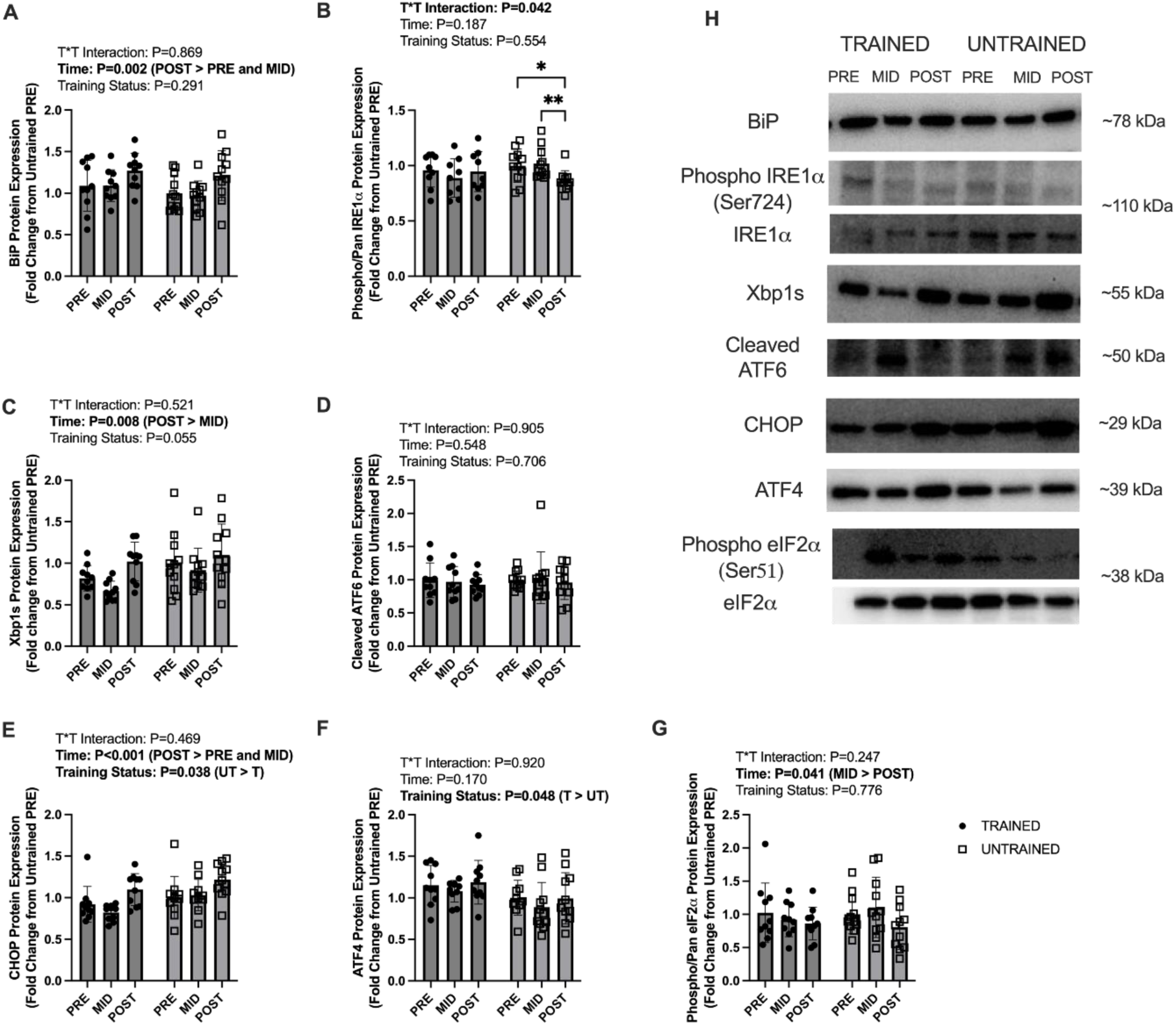
Endoplasmic reticulum stress responses. Legend: Data are presented as individual data points with mean and standard deviation bar graphs. Results from 21 human participants for BiP protein expression (panel a), phosphorylated/pan IRE1α (panel b), Xbp1s protein expression (panel c), cleaved ATF6 protein expression (panel d), CHOP protein expression (panel e), ATF4 protein expression (panel f), and phosphorylated/pan eIF2α protein expression (panel g). Panel h contains representative western blot images. Two-way ANOVA-derived P values for training status*time interactions, main effects of time, and main effects of training status are presented above graphs. For significant main effects, *post hoc* test details are presented in parentheses. *Post hoc* statistical significance between time points is presented for those ANOVAs demonstrating a significant training status*time interaction (panels a-d). Symbols: *, P<0.05; **, P<0.01. Abbreviations: BiP, binding immunoglobulin protein; IRE1α, inositol requiring enzyme 1 alpha; Xbp1s, spliced form of X-box binding protein 1; ATF6, activating transcription factor 6; CHOP, CCAAT enhancer binding protein (C/EBP) homologous protein; ATF4, activating transcription factor 4; eIF2α, eukaryotic initiation factor 2 alpha; phospho-, phosphorylated.

### Anabolic Markers

Phospho/Pan P70S6K1 did not demonstrate main effects of time or training status (P≥0.074) nor a training status*time interaction (P=0.473; Figure 8a). Phospho/Pan RPS6 also did not demonstrate main effects of time or training status (P≥0.178) nor a training status*time interaction (P=0.178; Figure 8b). Phospho/Pan 4EBP1 did not demonstrate main effects of time or training status (P≥0.269) nor a training status*time interaction (P=0.271; Figure 8c).

**Figure 8.**
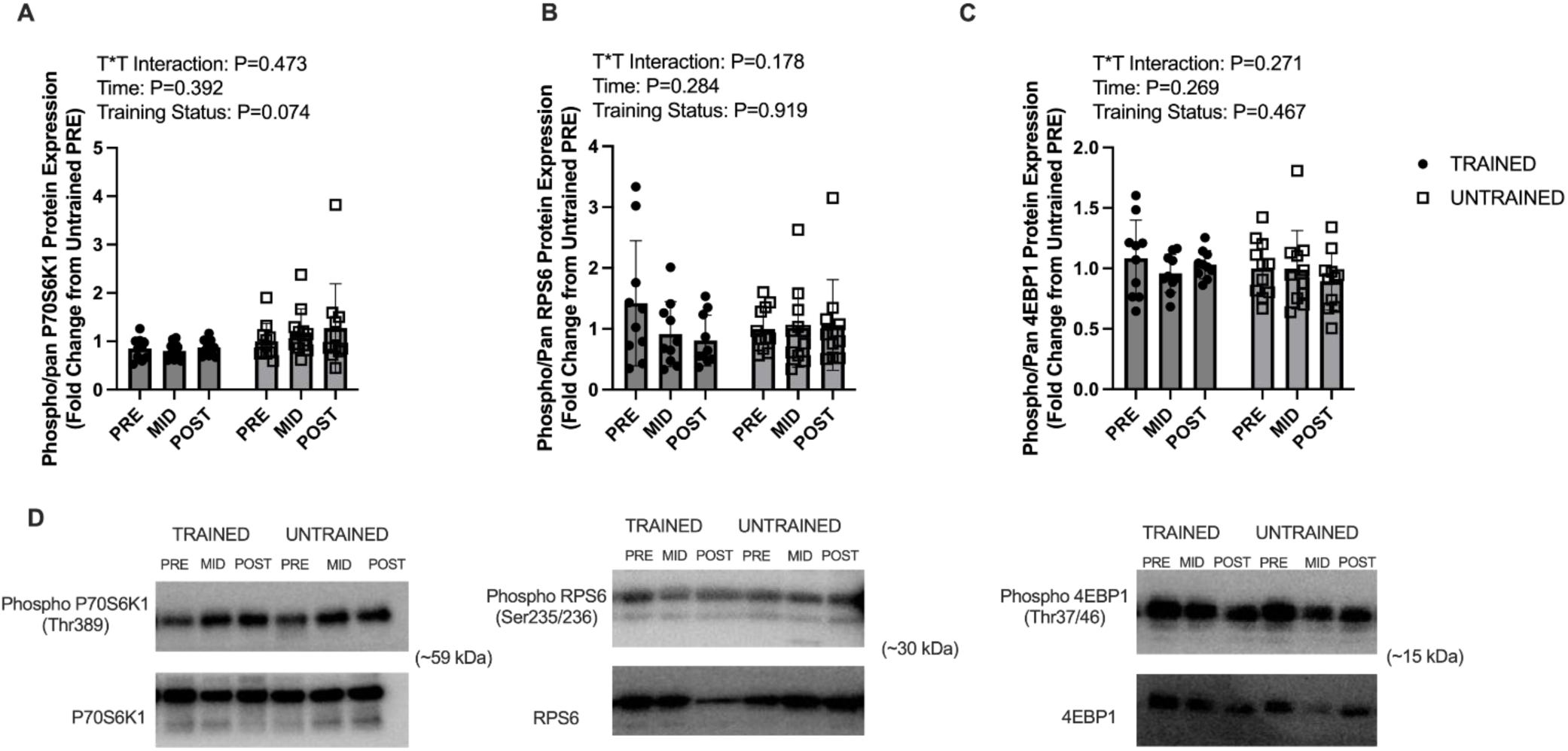
mTORC1 signaling responses. Legend: Data are presented as individual data points with standard deviation as error bars. Results from 21 human participants for phosphorylated/pan p70S6K1 protein expression (panel a), phosphorylated/pan rps61 (panel b), and phosphorylated/pan 4EBP1 (panel c). Panel d contains representative western blot images. Two-way ANOVA-derived P values for training status*time interactions, main effects of time, and main effects of training status are presented above graphs. Abbreviations: p70S6K1, ribosomal protein S6 kinase beta-1; rpS6, ribosomal protein S6; 4EBP1, eukaryotic translation initiation factor 4E-binding protein 1; phospho-, phosphorylated.

## DISCUSSION

The contributors to and effects of disuse in humans are somewhat characterized (6, 30). However, strategies to offset the maladaptation that accompanies limb immobilization are still emerging. The purpose of this study was to examine if RT status affects tissue and cellular-level adaptations to a future disuse event. We additionally examined if recovery adaptations from disuse, via eight weeks of prescribed/supervised RT, were differentially affected by training status. The primary finding herein was that both T and UT declined from PRE to MID in all macro-level measures of muscle size. Recovery RT led to UT exceeding their pre-intervention VL muscle size and T restoring their baseline muscle size. This response was generally recapitulated at the fiber level and was driven primarily by type II fiber plasticity. Additionally, both T and UT increased satellite cell number at POST, while T had greater satellite cell number and myonuclei per fiber across all time points. Total RNA content (a marker of ribosome content) declined with for T and UT, and 18S and 28S rRNA were responsive to RT, though primarily driven by the T group. Accompanying increases in rRNA content, two ribophagy proteins declined with RT, but the rRNA transcriptional drivers POLR1B and UBF paradoxically declined with RT. We also observed that the endoplasmic reticulum stress (ERS) proteins CHOP, BiP, and Xbp1s proteins were elevated with RT at POST for both groups, suggesting an elevated ERS response with RT as compared to leg immobilization. Finally, phosphorylation states of key anabolic signaling markers related to the mTORC1 pathway were not responsive to leg immobilization or RT. Each of these findings are discussed below in greater detail.

### Muscle Morphology

The primary finding from this study was that 14 days of leg immobilization predictably induced atrophy of the VL in both groups as determined through ultrasound. Moreover, it was observed through two different ultrasound measures (panoramic CSA; transverse thickness) that only UT exceeded baseline values of muscle size after eight weeks of RT. Notably, fCSA measures did not align completely with these ultrasound data, as mean fCSA was greater at POST and PRE than MID (main effect) while T was greater than UT throughout. This response was driven primarily by type II fibers. These discordant responses are not entirely surprising given that these measures do not recapitulate one another well. In this regard, data from our laboratory suggest that change scores in VL fCSA and ultrasound derived VL mCSA with RT do not significantly correlate (31). This could be due to heterogeneity of fiber type throughout the VL and potential differences in fiber type simply by biopsy site. Measurement differences notwithstanding, this intervention is the first to demonstrate differences in rates of recovery with RT after disuse in trained versus untrained individuals. Interestingly, only UT participants exceeded pre-intervention VL size values following the 8-week recovery RT program. Critically, T participants in this study had an extensive training history and it is possible that eight weeks of RT was not enough to produce significant hypertrophy above and beyond what these participants had accrued prior to the intervention. Seaborne et al. (22) reported that RT pre-conditioning is beneficial for VL size gain after a seven-week RT protocol, seven weeks of detraining (not disuse), and seven subsequent weeks of RT in untrained individuals. To this end, it is worth noting that the models utilized by each of these studies are markedly different; however, the notion of RT history eliciting a beneficial molecular backdrop (e.g. epigenetic memory) for future hypertrophy is possible. Furthermore, it is possible that, had this study followed the model set forth by Seaborne et al. (22), results could have been similar in T and UT. Given the current study design, it is not possible to parse out differences attributable to detraining (e.g. cessation of RT) versus leg immobilization.

While our data suggest that prior RT history does not mitigate muscle atrophy during a period of leg immobilization, it is difficult to surmise that there is no benefit of prior resistance training experience. For instance, both ultrasonography-based and fiber level data suggested that T maintained larger muscle size than UT throughout the intervention. Therefore, we maintain that habitual RT prior to disuse is likely beneficial by creating a physiological reserve (i.e. larger overall muscle size from which to lose) despite not affecting the rate of atrophy. It is also interesting that the observed fiber level changes were predominantly in type II fibers as opposed to type I fibers. Seminal work by the Booth laboratory suggests that the predominantly type I fiber-enriched soleus muscle in rodents is more susceptible to disuse atrophy than the type II fiber-enriched plantaris muscle (34). These data from the Booth laboratory are supported by certain findings in human spaceflight that show atrophy and maladaptation to myofilament spacing in the predominantly type I soleus muscle (35) along with findings suggesting that type I fibers of a given muscle preferentially atrophy as compared to the type II counterparts in that same muscle (36). It is however worth noting that Bass and colleagues (37) found that two different muscles of similar type I fiber predominance (tibialis anterior and medial gastrocnemius) demonstrated different susceptibilities to disuse-induced atrophy, with the medial gastrocnemius significantly declining in muscle volume and muscle thickness while the tibialis anterior remained unchanged. This is to say that in humans versus rodents it is possible that species differences in fiber type atrophy rate exist. Moreover, it is likely that divergent muscle-specific molecular factors play a critical role in determining the amount of atrophy that muscle will realize during a disuse period. To this end, it has been posited that the relative abundance of atrophy-protective genes (e.g. PGC1α) in type I fibers could have a protective effect in the absence of a stimulus such as denervation (rodent models only) and, thus, type II fibers could be more susceptible to muscle wasting induced by general disuse (reviewed in (38)). This remains speculative, as the protein expression of such genes were not measured herein, but the plasticity of type II fibers during the current intervention does shed light on the adaptations that occur to non-complicated immobilization-induced muscle atrophy and recovery RT in humans.

### Satellite Cell and Myonuclei Observations

Satellite cell number did not change with disuse but increased with RT. The observation that satellite cell number did not change with two weeks of immobilization suggests that the disuse protocol did not have notable effects on satellite cell proliferation and/or apoptosis. This observation has not been consistent between investigations or between models (reviewed (39)), as certain investigations have shown no change with bedrest or limb immobilization (40, 41), increases with immobilization (42), and decreases with spinal cord injury and aging (43). It should be noted that these findings are likely dependent on the disuse model employed and the age of participants undergoing disuse. Thus, it is possible that satellite cell number could be affected with a more severe and/or complicated disuse model or with an older cohort.

Contrary to the conflicting disuse model data, the response of satellite cells to mechanical overload is well-reported in both humans (reviewed in (4)) and rodents (reviewed in (44)). To this point, the increased satellite cell number with recovery RT align with this prior literature, and our finding that T possessed more myonuclei per fiber than UT is a well-supported phenomenon (45, 46). However, myonuclear number remaining unaffected from MID to POST in both groups is an interesting observation that warrants further discussion. As proposed by Murach et al. (44), several human studies have reported that skeletal muscle hypertrophy can occur with an increase in satellite cells and without myonuclear accretion. These authors additionally posit that satellite cells may adopt a fusion-independent role in supporting tissue remodeling through cell-cell communication via extracellular vesicles (EVs) (44, 47, 48). While aspects of this phenomenon were not assessed herein, we find this as a compelling possibility to explain our current observations; namely, the increase in satellite cell number with recovery RT could have been an adaptive response to aid in tissue remodeling via EV-mediated cell-cell communication. Follow up studies using high resolution molecular profiling such as long- and small-RNA sequencing would be useful efforts to assess whether differential expression of protein coding mRNA or analyses of mRNA targets of miRNA identify pathways or cellular processes involving satellite cells or EV trafficking.

### Ribosomal Markers

Markers of ribosome content (total RNA, 18S rRNA, 28S rRNA) appeared to decrease or not change with disuse and increase with recovery RT. In an investigation among middle aged men, Figueiredo et al. (49) reported a similar change with 14 days of leg disuse. In that investigation, these reductions with unloading were largely attributed to increases in RNA breakdown versus decrements in RNA synthesis (supported via mouse model). We measured surrogate markers suggestive of ribosome breakdown (i.e. ribophagy), and while these markers did not demonstrate a consistent response, ribophagy cannot be ruled out as a potential mechanism underlying the decrease in total RNA at MID. Further investigation utilizing tracers that can directly measure rates of synthesis and breakdown is needed to elucidate this phenomenon. The POL1 transcription factor UBF was reduced following RT, which appears discordant with findings suggesting ribosome biogenesis during the RT recovery phase. This was indeed unexpected. While we lack a clear explanation for this discrepancy, timing of the biopsy (at least 72 hours post-exercise) may have contributed. rDNA transcription is transient and precedes rRNA accumulation (reviewed in (50)); thus, it is likely that cellular UBF was not maintained in high protein concentrations after 8 weeks of RT. Interestingly, we observed an increase in POL1 protein expression at MID. This could be a compensatory mechanism by which myofibers attempted to overcome the lack of mechanical input (and, thus, lack of translation demand) by increasing ribosome content and maintaining overall translational output. Clearly, there is discordance by which ribosome content did not increase with elevated POL1 protein content. Hence, more investigation into the POL1 regulon with limb immobilization to elucidate this disconnect is warranted.

### Endoplasmic Reticulum Stress Proteins

The ERS and unfolded protein response (UPR) effectors BiP, Xbp1s, and CHOP were upregulated at POST as compared to MID (Xbp1s) or PRE and MID (BiP and CHOP). These three effectors of the UPR being upregulated at POST suggest participants in both groups had a larger ERS and/or unfolded protein burden after eight weeks of RT versus with two weeks of leg immobilization. In opposition to our findings, rodent and non-human primate data suggest ERS markers are upregulated with unloading (51, 52). Furthermore, it has been observed in younger adults that 9 days of bed rest upregulates ERS pathway representation via microarray analysis (53), and 8 weeks of RT in older adults reduces ERS pathway representation in peripheral blood mononuclear cells (54). Interestingly, however, Baehr et al. (55) reported that the ERS marker protein disulfide isomerase is increased with reloading in 9-month old rats subjected to 14 days of hindlimb unloading followed by 3, 7, or 14 days of reloading. This effect was amplified when 29-month-old rats were subjected to the same paradigm with effects extending to the ERS proteins CHOP and BiP. These rodent data somewhat align with our observations that RT-induced increases in muscle mass (but not unloading via leg immobilization) affect ERS markers. While speculative, we posit that the observed upregulation in these markers could be due to the robust hypertrophic response to RT (e.g. enhanced translational capacity/output, and more periods of positive net protein balance (56, 57)) rather than a mechanistic driver of hypertrophy. Nonetheless, research in this area has been primarily driven by rodent models and more human work is warranted.

### Anabolic Signaling Proteins

The phosphorylation statuses of key downstream mTORC1 effectors showed no differences between group or time. While it is often posited that decreased muscle protein synthesis is the key driver of disuse-induced atrophy (reviewed in (58)), it has been reported that the phosphorylation status of anabolic signaling proteins do not show remarkable variability with limb unloading. For instance, in a cohort of post-menopausal women undergoing disuse and rehabilitation (6 weeks of machine-based unilateral training), Moller et al. (59) reported no discernable pattern in changes to phosphorylation states of such proteins. Similarly, in a model of bedrest-induced atrophy in older humans preceded by either one unilateral RT bout or no training, Smeuninx et al. (60) reported no changes to these markers in the trained or untrained leg across time. In younger adults, Brook et al. (58) reported largely no differences in the phosphorylation states of mTORC1-related proteins in the control versus disuse legs following four days leg immobilization via bracing. Moreover, these observations coincided with marked atrophy and decrements in muscle protein synthesis. Our data are also confounded by the basal state at which all biopsies were taken in this study. Studies examining the responses of such proteins to mechanical overload typically show peak phosphorylation responses within a 24-hour period following a resistance exercise bout (61, 62, 63, 64, 65), while the biopsies taken herein occurred 72+ hours after the training bout in all cases. Hence, it remains possible that between-group differences in mTORC1 signaling could have occurred in the current study, albeit our sampling limitations precluded the ability to detect such differences.

### Limitations

This study is not without limitations, the primary being a relatively small sample size. Additionally, although recruitment efforts were aimed at having and equal sex-matched cohort of male and female participants, few female participants responded to our recruitment strategies to participate in the study. As such, sex-based comparisons were precluded, and many of the observed patterns herein were likely masked to some extent by a limited sample size in general. We are also limited by the lack of acute time point biopsies throughout training. While this would have been an insightful addition to the study, increasing participant burden was not desired. Finally, we are limited by the lack of direct metrics tracking adherence to the leg immobilization protocol (i.e. brace being removed at times not permitted or excessive loading when not permitted). Again, such measures would have provided additional insight; however, it is worth noting that the three different measures of muscle size suggest an overall adherence to the leg immobilization protocol.

### Conclusion

In previously trained and untrained younger adults, 14 days of leg immobilization produced similar VL muscle atrophy and 8 weeks of recovery RT induced similar hypertrophy outcomes. Moreover, molecular signatures between groups throughout the intervention were largely similar. Given that this study was limited to a younger adult population undergoing non-complicated disuse atrophy, more research is needed to determine if RT history elicits similar outcomes with aging or in individuals with medically complex limb immobilization issues.

## ADDITIONAL INFORMATION

### Author Contributions

JMM and MDR primarily drafted the manuscript; JMM and MDR prepared figures; HSB and MMV provided oversight during the disuse protocol; HSB served as the licensed physical therapist to assist with oversight throughout the intervention; MSS provided critical equipment and expertise for the carrying out of the leg immobilization protocol; JMM, JSG, CBM, and AAP primarily carried out laboratory based assays; ZAG and MMB provided critical resources for the measurement of ribosomal markers; DLP, MCM, MLM, PJA, BJM, DAA, and ACB provided crucial assistance with data collection procedures; DLP conducted all ultrasonographic imaging; MDR, JSG, CBM, HSB, ZAG, and MMB provided intellectual feedback for the duration of the project; All co-authors assisted with revising and editing the manuscript, and all co-authors approved the final version.

## ACKNOWLEDGEMENTS

We would like to acknowledge the participants who completed this study. We would like to acknowledge Jackson Dickey for his dedicated service in analyzing western blot images for this project.

